# Directed Human Structural Connectome Reveals Hierarchical Organization and Shapes Large-Scale Brain Dynamics

**DOI:** 10.64898/2026.06.16.732559

**Authors:** Nan Huang, Huifang Wang, Paul Triebkorn, Claudia Wheeler-Kingshott, Maciej Jedyank, Olivier David, Alain Destexhe, Egidio Ugo D’Angelo, Nigel Pedersen, Viktor Jirsa

## Abstract

The human structural connectome, most commonly derived from diffusion-weighted imaging (DWI) and tractography, provides a macroscopic description of whole-brain wiring and serves as the structural foundation of network neuroscience, large-scale brain simulations, and personalized digital brain twins. However, tractography-derived connectomes are fundamentally limited by their inability to distinguish afferent from efferent connections, yielding networks that are undirected and therefore blind to the hierarchical organization imposed by the directionality of anatomical connections. In this study, we introduce a directed human structural connectome (DHSC) by transferring tracer-derived projection patterns from macaque to human using cross-species connectivity blueprints. Topological analysis of the DHSC manifests biological plausibility, a small-world network organization, and a directionality-based hierarchy, which offer the hierarchical organization of human brain networks. In the context of brain dynamics, the introduction of directionality reshapes the propagation and persistence of sensory inputs. DHSC also best captures the empirical spatiotemporal dynamics of stimulus-evoked brain activity. The findings demonstrate that anatomical directionality is a critical determinant of large-scale brain organization and dynamics. This provides evidence that directed connectome may offer potential advantages in large-scale simulations of the human brain. The resulting DHSC, along with all related analyses and data are openly available.

## 1 Introduction

The human structural connectome, most commonly constructed using diffusion-weighted imaging (DWI) and tractography, provides a macroscopic description of whole-brain wiring [1–3], and has become a central substrate for network neuroscience [4, 5]. Diffusion MRI–based connectomes are now routinely used to study brain organization [4–6], disease-related network alterations [7, 8], and to parameterize whole-brain dynamical models [9, 10].

However, despite their widespread adoption, tractography-derived connectomes are subject to fundamental methodological limitations that constrain their anatomical and functional interpretability. Extensive validation studies comparing tractography with invasive neuroanatomical tracers have demonstrated that diffusion MRI tractography exhibits a systematic tradeoff between sensitivity and specificity, resulting in substantial rates of both false-positive and false-negative connections [11–15]. Tractography inherently limited in its ability to faithfully reconstruct long-distance projections and to resolve complex axonal trajectories, leading to distance-dependent biases [16]. Critically, diffusion MRI tractography also lacks sensitivity to axonal directionality. Because the diffusion signal reflects water displacement rather than synaptic or axonal transport direction, tractography cannot distinguish afferent from efferent connections and therefore yields connectomes that are effectively undirected or symmetric[17]. This limitation stands in sharp contrast to extensive neuroanatomical evidence from tract-tracing studies demonstrating that mammalian brain networks are intrinsically directed [18–20]. Directed connectivity is a fundamental organizing principle of brain circuits, shaping information flow, hierarchical processing, and the causal structure of large-scale dynamics. Directionality has been shown to distort network measures, alter hub organization, and obscure hierarchical structure [21]. From the perspective of the dynamic system, the introduction of asymmetry induces a shift from normal to non-normal[22], transforming degenerate invariant manifolds into structured flows with preferred trajectories [23]. Yet its consequences for whole-brain human dynamics have not been systematically characterized at the macroscale. Invasive neurotracing remains the gold standard for mapping axonal projections and their directionality, providing high-fidelity information about source–target relationships [18–20]. Although such methods cannot be applied in humans, the strong evolutionary conservation of large-scale brain organization between humans and non-human primates offers a principled opportunity to transfer anatomical knowledge across species [24–26].

Here, we leverage macaque neuro-tracing data [18] to infer a directed human structural connectome by transferring tracer-defined projection patterns to the human brain using cross-species connectivity blueprints [26]. By constraining diffusion MRI–based tractography with tracer-informed directionality, we derive a biologically grounded directed human structural connectome (DHSC). This approach allows us to augment conventional connectomes with probabilistic estimates of axonal polarity while preserving subject-specific tractography information. We confirmed the biological plausibility of the inferred connectome by comparing our results with the functional-anatomical connectome generated by the F-TRACT project. Then, we use this directed connectome to systematically characterize whole-brain network organization in the human brain, examining how directionality reshapes asymmetries, and hierarchical structure [27]. In particular, we quantify large-scale hierarchy using trophic-level organization, providing a directed, mechanistically interpretable measure of hierarchical positioning that complements existing functional and structural gradients [28]. Finally, we incorporate the DHSC into large-scale whole-brain dynamical simulations to directly test how biologically grounded directionality and altered topological features influence macroscale brain dynamics. By comparing directed and undirected connectomes under identical dynamics, we isolate the impact of directionality on resting-state and stimulus-evoked signal propagation dynamics. This framework provides a principled means to assess how incorporating anatomical directionality alters large-scale information flow and dynamical responses in the human brain.

Together, this work establishes a cross-species, tracer-informed strategy for constructing a directed human structural connectome and provides a foundation for evaluating the role of anatomical directionality in shaping large-scale human brain organization and dynamics.

## 2 Results

### 2.1 Constraining human tractography-based connectome via tracing based connection in Macaque

Neuroanatomical tracing provides precise maps of axonal projections but is inherently invasive, precluding whole-brain tracer-based connectomics in humans. We addressed this by leveraging evolutionary conservation between non-human primates and humans to transfer tracer-derived directionality from macaque to the human brain. By projecting macaque tracing data onto human brain maps, we informed our diffusion tractography with biologically grounded directionality. Specifically, we mapped macaque tracer connectivity [18, 19] onto the human brain using connectivity blueprints defined by 42 homologous white-matter tracts shared between the two species [26] (Fig. 1a). This approach establishes a cross-species correspondence between brain regions based on their tract-wise connectivity profiles, enabling principled transfer of projection patterns. Detailed procedures are provided in the Methods and Supplementary Fig. 1. Briefly, homologous white-matter tracts were reconstructed in macaque (Brainnetome-8 [12]) and human (100 HCP-YA subjects [29, 30]) datasets using tractography. For each cortical vertex and subcortical voxel, we computed its connectivity profile across the 42 tracts. These profiles were then averaged within each atlas region (macaque: D99 [31]; human: HCP-MMP1.0 [32] + HCPex [33]) to generate region-level connectivity blueprints (Fig. 1b). To infer human projections, the source and target regions of each macaque tracer-defined connection [18, 19] (CocoMac) were mapped to human regions based on the similarity of their blueprint profiles, quantified using symmetric Kullback–Leibler divergence (Fig. 1c). This procedure yielded, for each ordered pair of human brain regions, a probabilistic estimate of whether a connection exists. Since the original neurotracing matrix contained information only from the left hemisphere, we assumed identical connectivity patterns between the left and right hemispheres when inferring the right hemisphere. The differences in connectivity stem from the different match pattern in white matter structures between macaques and humans; for inter-hemispheric connections, we referred to the tracography structure (see the Methods section for detailed methodology). The resulting matrix, called directed human structural connectome (DHSC), provides a directed scaffold that informs the directionality of human tractography derived from macaque tract-tracing (Fig. 1a).

**Fig. 1.**
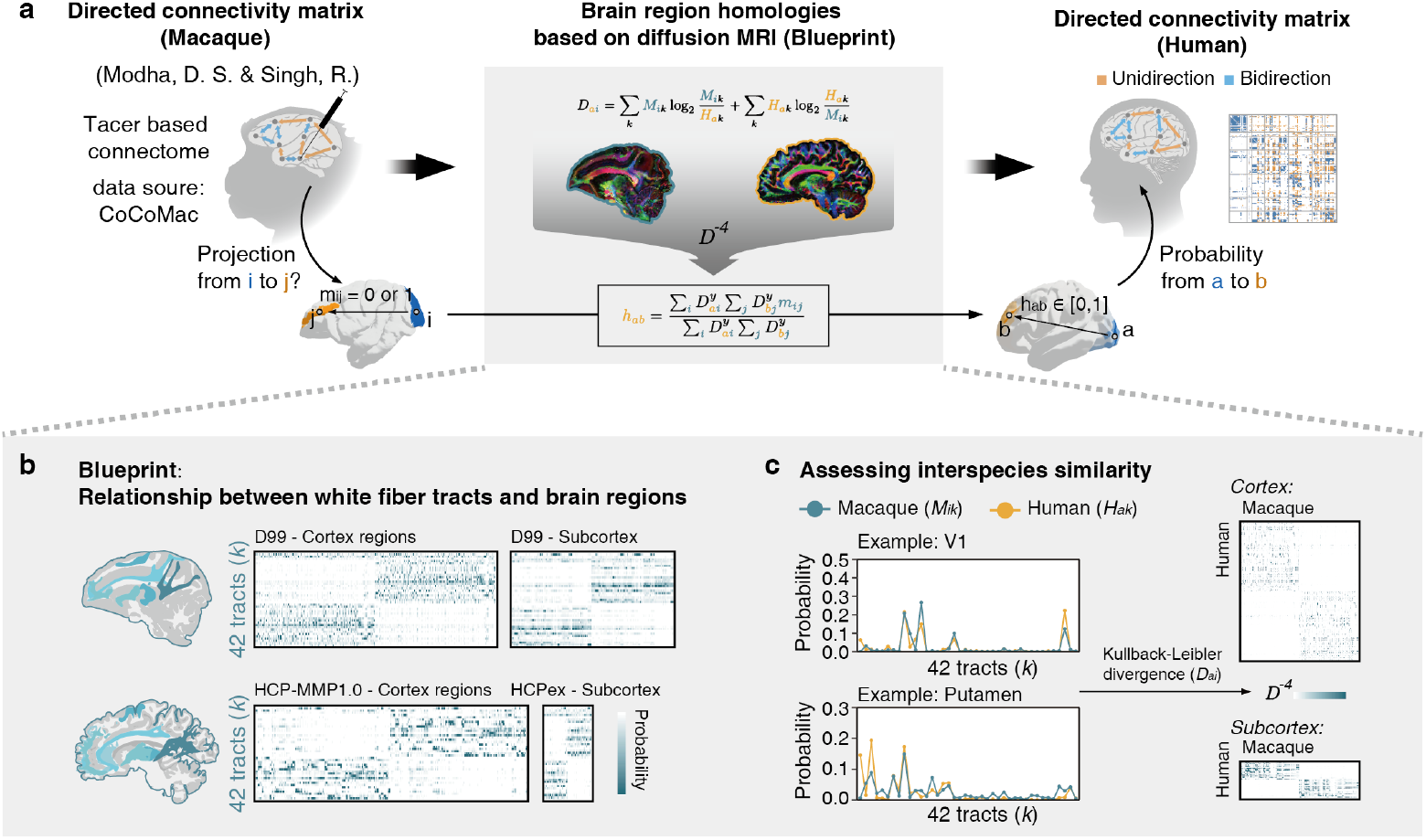
Inferring directed connectivity from macaque to human. **a**. Schematic illustrating the conceptual framework for mapping tracer based structural connectivity from macaque [18] to human. Owing to evolutionary conservation, 42 homologous white-matter tracts are shared across both species. By establishing region-to-tract connectivity relationships in the macaque [26], these relationships are subsequently transferred to the human brain to infer corresponding inter-regional connections (see Methods and Fig. S1 for details). **b**. Connectivity blueprints of macaque and human brains, representing region-wise projection profiles across the 42 homologous white-matter tracts. **c**. Example of cross-species connection mapping. Left: connectivity blueprints of two representative regions (V1 cortex and putamen) in macaque and human. Region similarity is quantified using the Kullback–Leibler divergence (*D*) between the distributions of tract-wise connectivity probabilities. Right: macaque-to-human mapping is performed by weighted average of divergence (*D*^*−*4^), enabling the transfer of directed connections onto the human atlas (HCP-MMP1.0 + HCPex).

To evaluate the biological plausibility of methodology, we compared the estimated connection probabilities against the cortico-cortical evoked potential (CCEP)–derived probabilistic effective connectivity matrix from the F-TRACT project in left cortex regions [34, 35], which demonstrates the effectiveness of this method under reasonable circumstances. CCEPs provide a direct, causal measure of effective cortico-cortical connectivity by quantifying stimulus-evoked responses that propagate along white matter pathways, with early components (N1) reflecting predominantly direct axonal projections. As such, F-TRACT offers an independent electrophysiological benchmark for large-scale human cortical connectivity and directionality. It is important to note that the connection probabilities estimated in the DHSC reflect the similarity of inferred projection origins and terminations, and therefore index the likelihood of anatomical connectivity rather than connectivity strength. Nevertheless, despite these conceptual differences, the DHSC and F-TRACT matrices exhibited substantial global structural similarity (Fig. 2a) between tracer-constrained tractography and stimulation-based effective connectivity. To balance biological plausibility with statistical robustness, we thresholded the DHSC, F-TRACT, and diffusion MRI (DWI) matrices to retain identical proportions of the strongest connections and performed density-dependent correlation analyses (Fig. 2b). The correlation between DHSC and F-TRACT peaked at a connection density of approximately 20%, a regime commonly considered to reflect intermediate network sparsity [4, 5] and to mitigate spurious weak connections while preserving large-scale organization (Fig. 2b). We therefore adopted a 20% density threshold for subsequent analyses. At this density, the overall network architectures of DHSC and F-TRACT were highly concordant, with discrepancies primarily confined to partially connected edges (Fig. 2c). Notably, relative to conventional DWI-based connectomes, DHSC preserved a greater proportion of long-range cortico-cortical connections (Fig. 2d), consistent with extensive tracer evidence that long-range association fibers are systematically under-represented in diffusion tractography [11, 12]. In addition, DHSC exhibited increased network asymmetry (Fig. 2e,f), in line with neuroanatomical tracer studies demonstrating pervasive directionality and hierarchical organization of cortical projections [36]. Finally, we observed spatially structured changes in nodal in- and out-degree, including increased out-degree in occipital-parietal lobe coupled with elevated in-degree in frontal regions (Fig. 2g). This pattern is consistent with known feedforward–feedback organization and long-range occipito-frontal pathways supporting visual–cognitive integration [37, 38], further supporting the biological plausibility of the DHSC architecture.

**Fig. 2.**
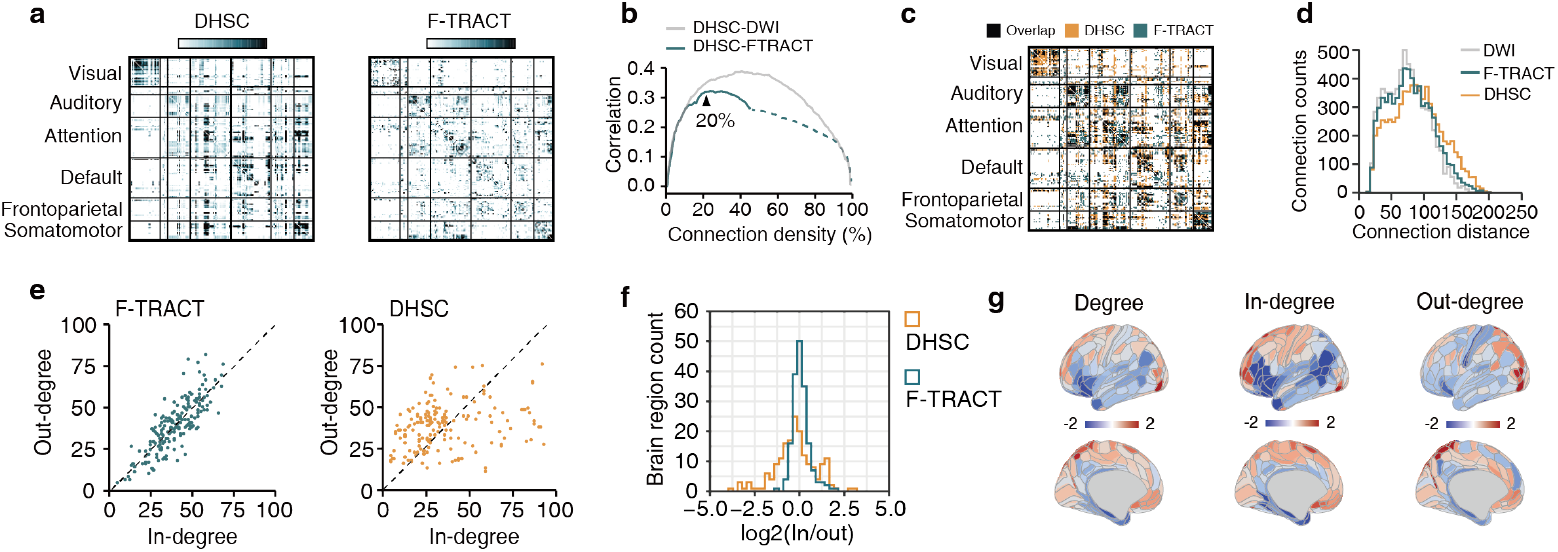
Cross-validation and comparison with a human directed connectome derived from cortico-cortical evoked potentials (CCEP) **a**. Putative cortico-cortical connections in the left cerebral hemisphere. Left: inferred directed human structural connectome (DHSC) derived from macaque tracer data. Right: human directed connectome derived from cortico-cortical evoked potentials (CCEP) from the F-TRACT project. Color represents the possibility of connection: white, low connection possibility; Dark-blue, high connection possibility. [35]. **b**. Correlation analysis between the F-TRACT connectivity matrix, the DHSC, and the DWI-based connectivity matrix. All matrices were thresholded to retain the same connection density and binarized prior to comparison. Gray: correlation between DHSC and DWI; Blue: correlation between DHSC and F-TRACT, Since FTRACT does not detect all possible connections, dashed lines are used to indicate connections that extend beyond the detection boundaries (The matrix remains unchanged after going out of bounds). **c**. Edge-wise overlap between the F-TRACT connectome and the DHSC at 20% connection density. Black: shared connections; orange: DHSC-only connections; blue: F-TRACT-only connections. **d**. Distribution of connection lengths (streamline). Grey: DWI-based connections; orange: DHSC connections; blue: F-TRACT connections. **e**. Regional in-degree and out-degree distributions. Left: F-TRACT connectome; right: DHSC. **f**. Distributions of the in-/out-degree ratio for DHSC and F-TRACT. orange: DHSC; blue: F-TRACT. **g**. Spatial distribution of the degree difference between DHSC and F-TRACT connectome, including also seperately indegree and outdegree. The degree difference *dd* is defined as *log*_2_(*Degree*_*DHSC*_*/Degree*_*F −T RACT*_ ). If both connectomes have the same degree, *dd* = 0; if DHSC has the higher degree, *dd* > 0; otherwise, *dd* < 0.

### 2.2 Basic network architecture of directed human structural connectome (DHSC)

DHSC identified 9,121 directed edges among each hemisphere brain regions (360 cortical regions from HCP-MMP1.0 [32] and 66 subcortical regions [33] from HCPex) of which 24.7% were unidirectional (Fig. 3a). We further quantified the regional distribution of bidirectional versus unidirectional connectivity. Visual cortical areas were enriched for bidirectional connections, consistent with dense reciprocal cortico-cortical interactions supporting hierarchical and recurrent visual processing [39]. In contrast, the insula, parietal-occipital regions exhibited a higher proportion of unidirectional connections, suggesting more asymmetric projection patterns of posterior parietal cortex circuitry and insula [40, 41] (Fig. 3a). At the whole-brain level, out-degree—but not in-degree—showed a significant positive correlation with task-positive scores derived from HCP task-fMRI [32] (r = 0.431, *P* < 0.001; Fig. 3b,c). This result offers indirect evidence for the biological plausibility of DHSC, while also suggest that certain nodes, spreading information broadly, are the key driver of how the brain activity changes [42, 43]. To characterize large-scale network organization, each region was annotated according to canonical functional systems defined by Ji et al [44], based on resting-state fMRI network partitioning (Fig. 3d, left). Consistent with extensive prior work showing that both structural and functional brain networks exhibit small-world topology, we quantified small-worldness using the *ω* metric proposed by Telesford et al. [45] The DHSC network exhibited small-world organization with *ω* = −0.088 (where *ω* = 0 indicates an optimal small-world regime), reflecting high clustering combined with relatively short path lengths. This architecture supports efficient segregation and integration, a hallmark of biological brain networks. At the mesoscale, functionally related regions formed clustered subnetworks, interconnected by a subset of long-range directed links that act as shortcuts between modules (Fig. 3d). This pattern is consistent with known principles of cortical organization, in which long-range projections enable rapid integration across distributed functional systems. At the circuit level, DHSC recapitulated several well-established projection motifs. Primary visual cortex (V1) formed reciprocal connections with higher-order visual areas, consistent with known feedforward–feedback hierarchies in visual cortex [46]. Area 4 (primary motor cortex) integrated parietal inputs and exhibited reciprocal connectivity with premotor cortex, consistent with sensorimotor interactions [47]. The superior temporal visual area (STV) received projections from auditory cortex and projected to prefrontal cortex, in line with multisensory integration and auditory–prefrontal pathways [48]. Area 9p received widespread cortical inputs and showed reciprocal connectivity with prefrontal regions, consistent with its role in higher-order executive and associative processing [49] (Fig. 3e). Finally, DHSC captured the canonical cortico–basal ganglia–thalamo–cortical loop, including projections from prefrontal cortex to basal ganglia, from basal ganglia to thalamus, and from thalamus back to cortex [50] (Fig. 3f). The predominantly unidirectional nature of cortical-to-basal ganglia projections and the structured relay through thalamus are well-established features of these circuits and are critical for action selection, cognitive control, and reinforcement learning. The recovery of these motifs further supports the biological plausibility of the directed human structural connectome.

**Fig. 3.**
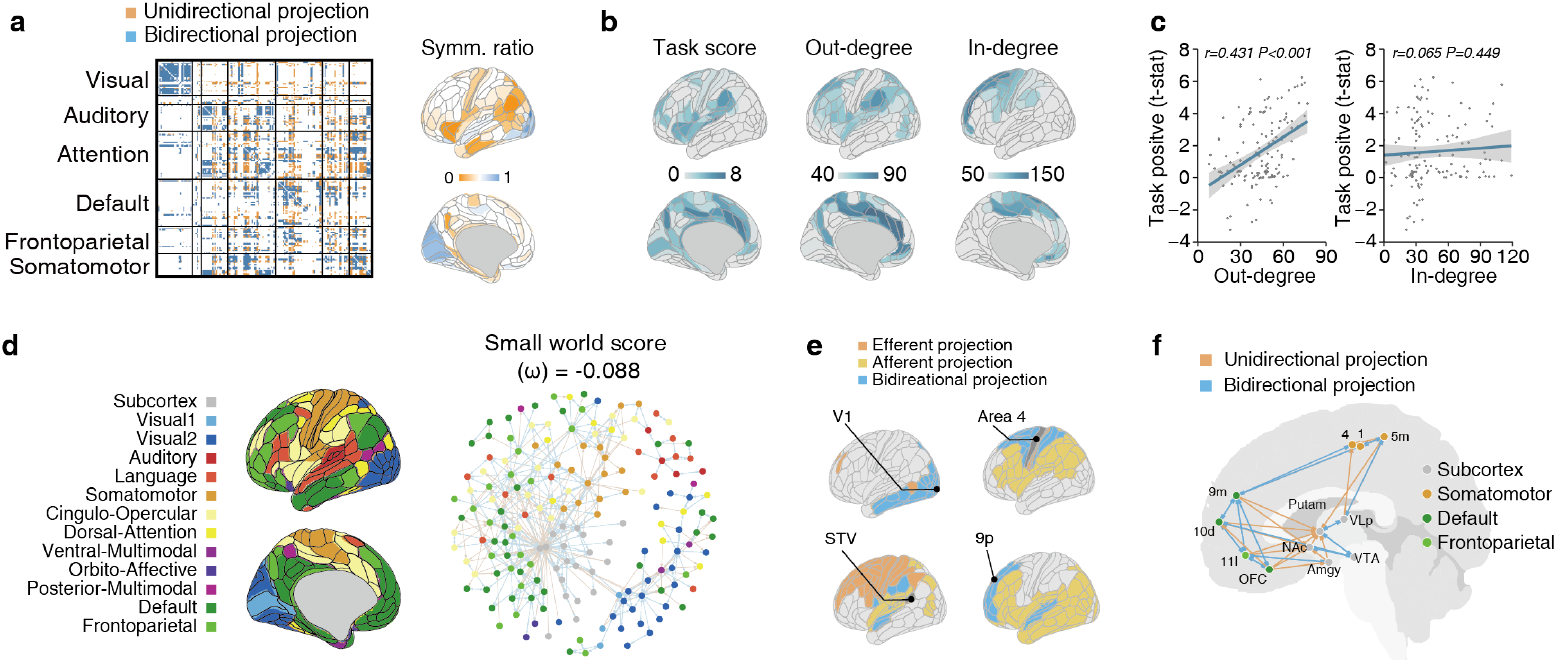
Basic topological characteristics of the directed human structural connectome (DHSC) **a**. Asymmetric connectivity in the DHSC. Left: connectivity matrix highlighting unidirectional (orange) and bidirectional (blue) connections between brain regions. Right: regional symmetry ratio projected onto the cortical surface. **b**. Spatial correspondence between task-positive regions (task-positive scores comes from Human connectome project[32]) and nodal out-degree. From left to right: cortical task-positive scores, out-degree distribution, and in-degree distribution. **c**. Correlations between task-positive scores and nodal out-degree and in-degree in the DHSC. **d**. Small-world organization of the DHSC. Left: canonical functional network parcellation of the cerebral cortex derived from resting-state fMRI [44]. Right: whole-brain directed connectivity network (top 500 strongest connections), with bidirectional edges shown in light blue and unidirectional edges in orange. Node colors denote functional network affiliation. The small-world coefficient (*ω*) is indicated above the network visualization. **e**. Directed connectivity patterns of representative regions (V1, area 1, STV, and 9p). **f**. The directed connectome reveals a canonical cortico–subcortical–cortical circuit motif, involving prefrontal cortex, striatum, thalamus, and cortex.

### 2.3 Hierarchical organization of the directed human structural connectome (DHSC)

The hierarchical organization of the brain is a central principle of large-scale cortical and subcortical architecture. Numerous landmark studies have characterized this hierarchy using diverse modalities, structural and functional connectivity gradients [28], myelin [51, 52], and intrinsic timescale gradients [53–55]. Together, these approaches consistently reveal a macroscale hierarchy spanning from unimodal sensory regions to transmodal association cortex, particularly default mode and frontoparietal systems, supporting increasingly integrative and abstract processing. While these prior approaches have provided convergent evidence for hierarchical gradients, directed connectivity networks in human offer a complementary and mechanistically grounded framework, as they explicitly encode the direction of information flow. This enables direct estimation of hierarchical ordering based on asymmetric projections. Accordingly, we adopted the concept of trophic levels from ecological network theory [27] to quantify hierarchical structure in the directed human structural connectome. In this framework, analogous to energy flow in food webs, information is modeled as flowing from lower- to higher-level regions. We further quantified trophic incoherence (*F*_0_) to assess the extent to which network organization adheres to a coherent hierarchical flow, with lower *F*_0_ indicating stronger hierarchical consistency [27] (Fig. 4a).

**Fig. 4.**
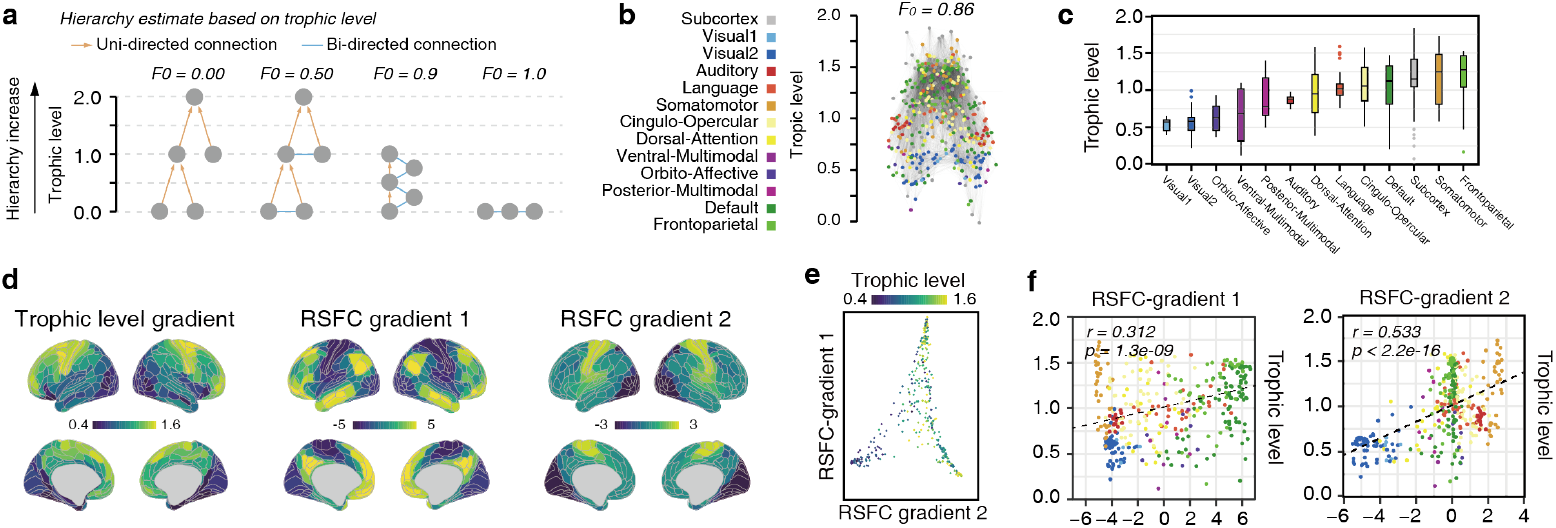
Hierarchical brain network organization inferred from the directed human structural connectome (DHSC) **a**.Framework for quantifying hierarchical organization from directed connectivity [27]. The schematic illustrates the continuum from a perfectly hierarchical network (trophic incoherence *F*_0_ = 0) to a fully non-hierarchical network (*F*_0_ = 1). Node-specific trophic levels estimate hierarchical position, while trophic incoherence (*F*_0_) quantifies the global degree of hierarchy in the network. **b**.Whole-brain hierarchical organization derived from the DHSC. The vertical axis denotes trophic levels, and node colors represent functional brain networks defined by resting-state fMRI. **c**.Distribution of trophic levels across functional networks. **d**.Cortical surface maps of the trophic-level gradient and the resting-state functional connectivity (RSFC) gradient [28]. **e**.Relationship between the trophic-level gradient and RSFC gradients 1 and 2. Point color indicates trophic level. **f**.Correlations between regional trophic levels and RSFC gradients 1 and 2. Point colors denote functional network affiliation, consistent with panel b.

Analysis of the DHSC revealed a pronounced and biologically interpretable hierarchy. Visual and auditory networks occupied lower trophic levels, whereas the default mode network, frontoparietal control network, and motor cortex exhibited higher trophic levels (Fig. 4b,c). This ordering is consistent with extensive evidence placing primary sensory cortices at the lower end of cortical hierarchies and transmodal association systems at higher levels of integration. Despite the network displaying a clear hierarchical structure, the addition of numerous unidirectional edges does not result in a purely feedforward architecture (*F*_0_ = 0.86). Instead, reciprocal and looped motifs dominate the network. We next compared trophic level gradients with canonical resting-state functional connectivity gradients [28] (Fig. 4d-f). The principal rs-fMRI gradient (RSFC-gradient 1), commonly interpreted as a unimodal-to-transmodal hierarchical axis, showed only a modest correlation with trophic level (r = 0.312). In contrast, trophic level exhibited a stronger correlation with RSFC-gradient 2 (r = 0.533) (Fig. 4f).

### 2.4 Impact of directed connectome topology on whole-brain dynamics

Understanding how structural network architecture shapes large-scale brain dynamics is a central problem in systems neuroscience. In particular, it remains unclear to what extent topological features such as long-range connectivity and connection directionality influence emergent whole-brain dynamics. To address this question, we employed The Virtual Brain (TVB) platform to perform large-scale simulations, systematically testing how different connectome representations affect macroscopic neural dynamics (Fig. 5a, Fig. S3). Each brain region was modeled as a neural mass node with identical intrinsic dynamics, based on the mean-field reduction of a fully connected network of quadratic integrate-and-fire (QIF) neurons [56]. This approach limits the number of free parameters by avoiding region-specific parameter optimization, thereby enabling direct attribution of dynamical differences to connectome structure rather than local model heterogeneity. To isolate the effects of connectome topology and directionality, we adopted a parsimonious modeling strategy. We simulated whole-brain dynamics using three distinct connectome representations: (1) a diffusion MRI–based structural connectome (DWI), widely used in large-scale human brain modeling; (2) a symmetrized version of the directed human structural connectome (s-DHSC), used to keep topological property of DHSC while removing directionality; and (3) our proposed DHSC. All neuroimaging data used for simulation were drawn from the HCP-YA dataset [29] (n = 100; see Methods).

**Fig. 5.**
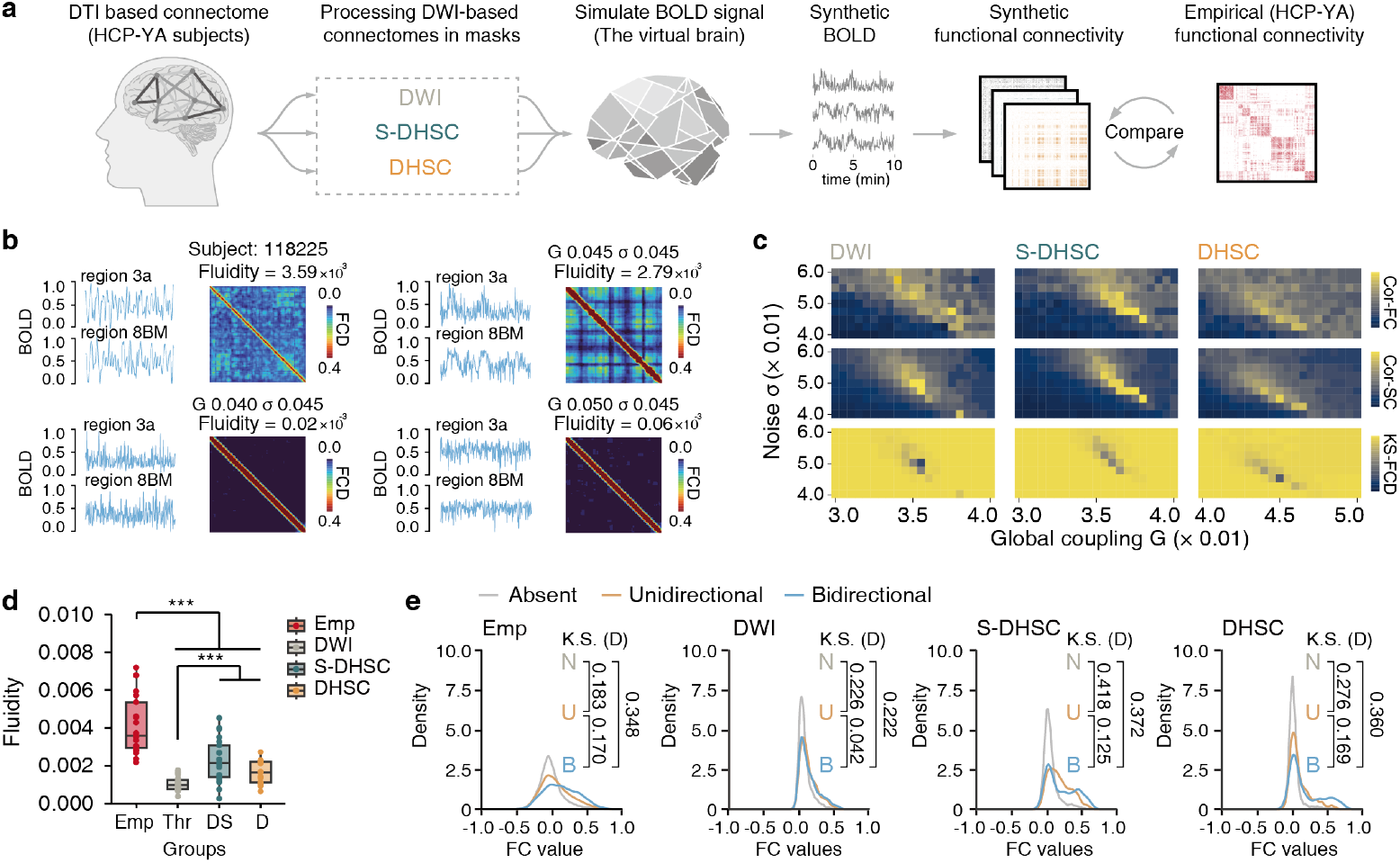
Effects of directed structural topology on resting-state whole-brain dynamics **a**. Overview of whole-brain simulation framework. Twenty subjects were randomly selected from the HCP-YA dataset. For each subject, three structural connectomes were constructed: (1) DWI: undirected connectome based on streamline counts, thresholded to the top 20 % of connections; (2) Symmetric-DHSC (S-DHSC): DWI connectome masked by the symmetrized directed scaffold; (3) DHSC: DWI connectome masked by the inferred directed human scaffold. Each connectome was implemented in The Virtual Brain platform (see Methods and Fig. S3). Simulated BOLD functional connectivity (FC) was compared with empirical resting-state FC from the corresponding subjects. **b**. The examples illustrate the FCD for empirical data and simulated data, respectively. The top-left figure shows empirical data; the rest are simulation results for different values of global coupling strength (*G*) under same noise (*σ* = 0.045). **c**. Model control parameters and goodness-of-fit metrics. *G* and *σ* served as control parameters. Model performance was evaluated using three criteria: (i) correlation between simulated and empirical FC, (ii) correlation between simulated FC and empirical SC, and (iii) similarity between simulated and empirical functional connectivity dynamics (FCD) distributions. **d**. Individual-level comparison of simulated and empirical fluidity. Distributions are shown for each connectome type. ***: t.test indicates *P <* 0.001. **e**. The relationship between structural and functional connections under experimental and simulated conditions. Brain regions were divided into three groups based on DHSC: no connections, unidirectional connections, and bidirectional connections. The density map illustrates the distribution of functional connectivity across different structural connections (from left to right: empirical data, and simulation results based on DWI, S-DHSC, and DHSC). Kolmogorov–Smirnov test (K.S.) were used to quantify the distribution difference, the distribution distance (D) is presented in the figure.

Criticality denotes a dynamical regime near the boundary between ordered and disordered states [57]. In brain networks, critical dynamics are hypothesized to optimize information processing by balancing stability and flexibility: in the subcritical regime, activity is stable but signal propagation decays; in the supercritical regime, activity persists but becomes less informative (Fig. 5b). Therefore, for each connectome, we tuned the global coupling parameter (*G*) which modulates inter-node interactions and the noise amplitude (*σ*) which controls spontaneous activity to bringing the system toward a critical regime that we treated as the resting operating point (Fig. 5c). To characterize this regime, we used three metrics: (1) the correlation between simulated and empirical functional connectivity; (2) the correlation between simulated functional connectivity and structural connectivity; and (3) the similarity between simulated and empirical FCD matrix distributions, assessed with the Kolmogorov–Smirnov test. The parameter combinations with highest Fluidity value and lowest KS value were selected as resting operating point (DWI: *G* = 0.0355, *σ* = 0.050; S-DHSC: *G* = 0.0360, *σ* = 0.050; DHSC: *G* = 0.0450, *σ* = 0.045) (Fig. 5c).

To test whether reshaping of the connectome altered resting-state dynamics, we reshaped the connectomes of 20 HCP-YA subjects and performed independent parameter sweeps. We quantified fluidity using the variance of the FCD matrix. To obtain a single connectome-level fluidity score, we then took the maximum fluidity observed across the parameter space. The results showed that substantial topological pruning (S-DHSC) significantly increased resting-state fluidity, whereas adding only directionality (DHSC) did not produce a significant change in fluidity (Fig. 5d). We then examined whether structural differences were reflected in static functional connectivity. Using DHSC connectivity patterns, we classified edges into three categories: absent, unidirectional, and bidirectional. In the empirical data, unidirectional and bidirectional connections showed a graded separation (Fig. 5e, left), supporting the plausibility of the DHSC. In contrast, the DWI control network showed no distinction between unidirectional and bidirectional categories. Both the s-DHSC and DHSC networks exhibited such distinctions, with the DHSC showing a stronger gradient (Fig. 5e). Overall, pruning of the connectome can alter resting-state dynamics, whereas the addition of directionality has a comparatively limited effect within the present modeling framework, a whole-brain simulation whose nodes are modeled as QIF-based mean-field neural masses.

### 2.5 Directionality reshapes stimulus-evoked signal propagation

Finally, building on parameter regimes established under resting-state conditions, we examined whether connectome topology and directionality shape stimulus-evoked signal propagation at the whole-brain scale. Using the same three connectome representations (DWI, s-DHSC, and DHSC), we applied a 2 s square-pulse input to primary visual (V1) and primary auditory (A1) cortices to approximate transient sensory drive. Stimulus-evoked activity was obtained by subtracting baseline dynamics, and propagation was characterized using region-wise time series and waveform features (Fig. 6a).

**Fig. 6.**
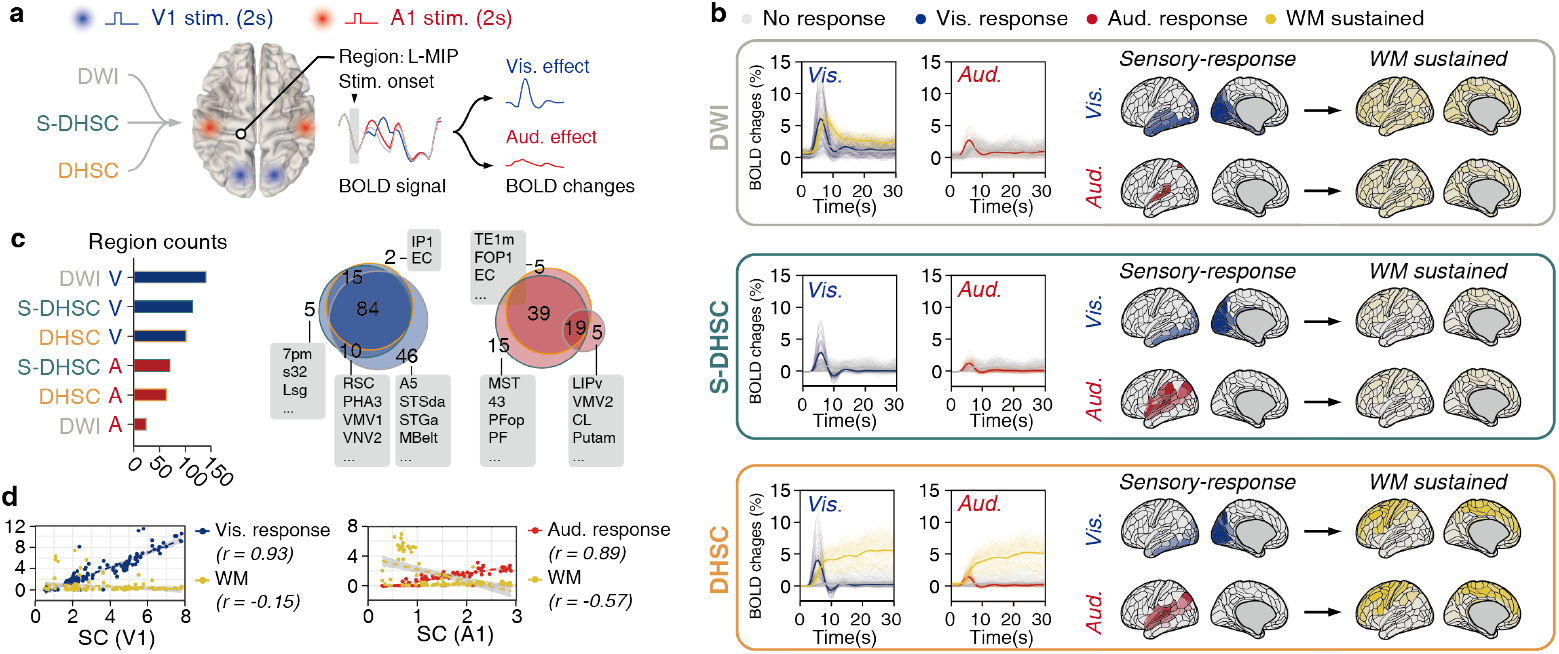
Directionality shapes whole brain response to sensory stimulus **a**. Overview of the stimulus-driven whole-brain simulation framework. Using resting-state regimes derived from three connectomes (DWI, S-DHSC, and DHSC), a independent 2-second external square-pulse stimulus was applied to primary visual cortex (L-V1, R-V1) and primary auditory cortex (L-A1, R-A1) to simulate visual and auditory inputs. By subtracting baseline activity (resting oscillation) from the perturbed state, stimulus-evoked propagation time series were extracted across all brain regions. **b**. Profile of whole brain dynamics constrained by different connectomes, The three panels, from top to bottom, show the simulation results based on DWI, S-DHSC, and DHSC, respectively. Left panel: Simulated time series induced by visual and auditory stimulus, three typical response patterns were identified: 1. sensory response to stimuli in the initial phase (blue: visual response; red: auditory response); 2. Working memory sustain (yellow, average BOLD change > 2% during 20− 30 s after stimulus onset); 3.no response (gray). Each thin line represents the average response of a single brain region to the input stimulus, with the color indicating the corresponding pattern. The thick lines represent the average activity of brain regions exhibiting the same response pattern. Right panel shows the distributions of sensory response (maxium BOLD changes in 0-10 seconds after stimulus onset) across the whole brain. **c**. The overlap between brain regions activated by different connectomes and external stimulus. The bar chart on the left shows the total counts of brain region activations under different conditions. Red represents auditory input signals, and blue represents visual input signals. The intensity of the color indicates the peak amplitude of the signal. The Venn diagram on the right illustrates the overlap between different sets, and the primary brain regions where the sets differ are listed. The filled color represents the stimulus type. (blue: visual stimulus, red: auditory stimulus); The outline represent the connectome used for simulation. **d**. The relationship between auditory and visual response, working memory sustained, and structural connections to brain regions receiving stimulus (V1, A1) using DHSC.

In contrast to resting-state results, stimulus-evoked dynamics were sensitive to directionality. We categorize regional responses based on post-stimulus activity and amplitude into two distinct response behavior: a rapidly decaying response with weak sustained activity (Fig. 6b, vistual: blue, auditory: red), and prolonged post-stimulus activity (Fig. 6b, yellow). We defined working memory–like maintenance [58, 59] as sustained activation over a 20–30 s post-stimulus interval. Under this definition, only the DHSC connectome supported robust, modality-independent sustained activity, whereas this property was largely abolished after symmetrization (s-DHSC) (Fig. 6b). We then compared the effects of connectome directionality on whole-brain dynamics, focusing on the initial and maintenance phases of stimulus propagation.

Regarding the initial stage of stimulus-response propagation, connectivity scaffold altered the spatial extent of rapid responses, restricting the spread of visual input to dorsal temporal regions (e.g., A5, STSda, MBelt) and reducing bias between visual and auditory pathways (Fig. 6b, c). Across DHSC and s-DHSC, regions exhibiting rapid responses largely overlapped (visual: 99/116, auditory: 58/78), but directionality introduced subtle spatial refinements, including less selective recruitment of regions such as retrosplenial cortex (RSC) for visual input and inferior parietal regions (PF, PFop) for auditory input, improving qualitative agreement with known sensory processing pathways [60] (Fig. 6b, c, Fig. S5). Consistent with network communication principles, early response amplitude was strongly associated with direct structural connectivity to the stimulated regions (visual: *r* = 0.93, auditory: *r* = 0.89), whereas sustained activity showed a weaker or negative association with these direct inputs (visual: *r* = −0.15, auditory: *r* = −0.57) (Fig. 6d), indicating that introducing directionality promotes transition from feedforward propagation to distributed recurrent dynamics [4, 5]. In other words, feedforward propagation refers to signals flowing outward from the source along direct connections during the early phase. During the sustained phase, signals instead circulate across many regions through feedback loops (Fig. 7b, e).

**Fig. 7.**
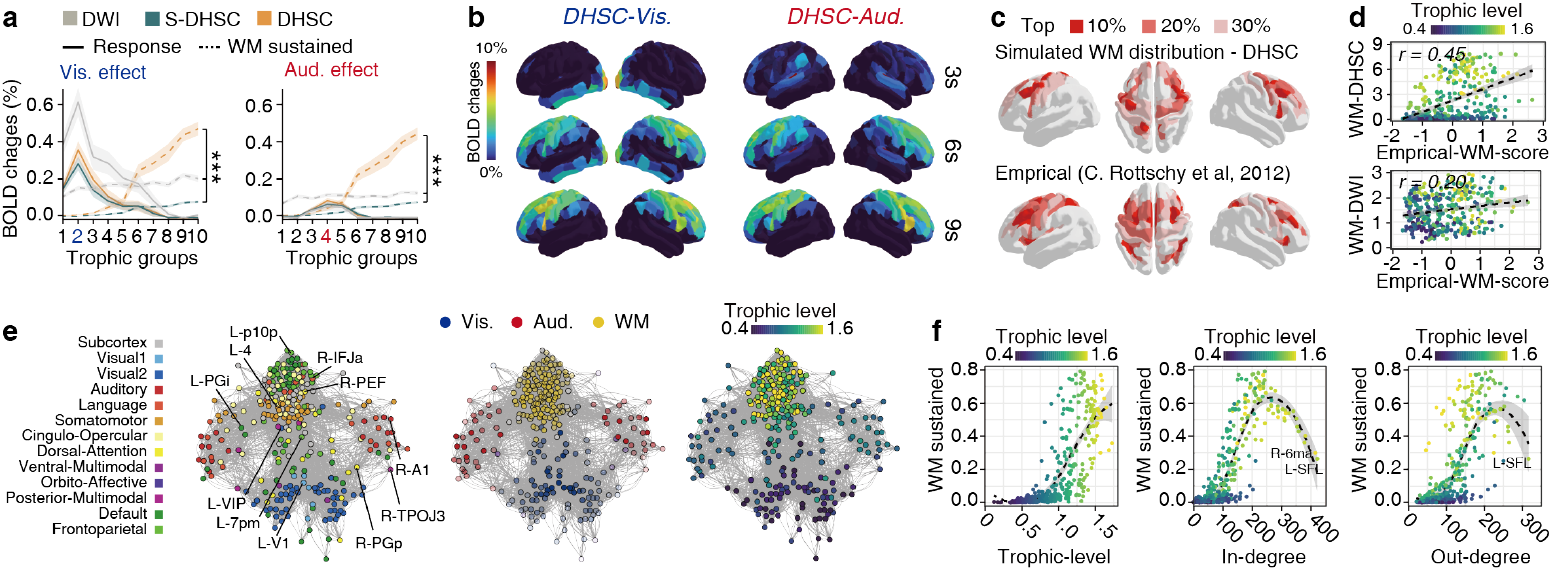
Directionality based hierarchy contribute to the formation and sustenance of working memory **a**. The relationship between auditory and visual response, working memory sustained, and trophic levels. The colors indicate the connection groups used for the simulation. The solid line and the dashed line represent the response to stimulus and the maintenance of working memory. Each point represents a brain region; only those regions directly connected to V1 (left) or A1 (right) are shown in the figure. **b**. Propagation of responses triggered by external stimuli under DHSC constraints. The colors represent changes in the BOLD signal. **c**.The relationship between simulation-based and experiment-based working memory distributions. The distribution of experiment-based working memory and simulation-based working memory in whole brain (showing the top 30% of brain regions). **d**. The correlation between working memory distributions based on DHSC and DWI and those based on experimental data. Each point represents a brain region and color represent the trophic level. **e**. The node network diagram illustrates visual and auditory signals converge within the same network to form working memory in DHSC based simulation. Left: color represent the RS-functional network, middle: color represent the response pattern, right: color represent trophic level. **f**. The relationship between working memory signals and structural connectivity network features based on DHSC simulation. The left, middle, and right panels illustrate the relationships between working memory and trophic levels, as well as in-degree and out-degree.

To further characterize signal propagation and maintenance in whole brain, we quantify the response to sensory stimulus as the maximum bold changes in fast response phase (0-10s after stimulus onset) and working memory as mean bold change during sustentation phase (20-30s after stimulus onset), and map to trophic hierarchy groups (Fig. 4b, c, d). During the early response phase, all connectome variants exhibited similar hierarchical decay of activity (Fig. 7a). In contrast, during the maintenance phase, marked differences emerged (anova-test, visual: *P* < 0.001, auditory: *P <* 0.001): in DWI-based simulations, sustained activity was weak and broadly distributed across all trophic levels (Fig. 6b, 7a), whereas in DHSC-based simulations, it was preferentially concentrated in association regions, particularly in parietal and prefrontal cortices (Fig. 7b). Notably, sustained brain activity is highly correlated between visual and auditory stimuli (*r* = 0.99, Fig. 6b, 7b), consistent with the hypothesis that information from different modalities is integrated in associated brain regions. This dynamic feature was largely disrupted after symmetrization of the connectome (Fig. 6b). We further compared the spatial distribution of sustained activity with empirical meta-analytic maps of working memory [61]. DHSC-based simulations showed stronger correspondence with empirical patterns (*r* = 0.45), particularly in prefrontal regions (e.g., Brodmann areas BA6, BA8, BA24, and IFJ), whereas DWI-based simulations exhibited weaker and more diffuse correspondence (Fig. 7c, d). Furthermore, sustained activity was consistent with the hierarchy predicted by trophic level due to the directionality, yet was not trivially explained by local network properties such as in-degree or out-degree (Fig. 7f), some structurally central nodes (e.g., 6ma, SFL) did not exhibit strong maintenance-related activity (Fig. 7f).

We also demonstrated that hierarchical network architecture drives form directional connectivity shapes the balance between rapid sensory processing in unimodal regions (sensory regions in blue and red) and sustained integration in transmodal cortical systems (association region in yellow) (Fig. 7e). In summary, our DHSC incorporating connectome topology and directionality exerts a dominant influence on how stimulus-evoked activity spreads and is sustained across the brains, and better matches empirical observations.

## 3 Discussion

### 3.1 A principled framework for inferring the DHSC

This study introduces a principled framework to infer a DHSC by transferring tracer-derived projection patterns from macaque to human via white-matter connectivity blueprints. The blueprint approach provides a biologically grounded mapping between homologous tracts across species and permits the principled projection of mesoscale tracer information onto human parcellations without direct invasive tracing in humans (Fig. 1a). The approach addresses a core limitation of diffusion MRI, although tractography provides valuable macroscopic wiring estimates, it systematically suffers from false positives, false negatives and ambiguity about axonal polarity, which constrain its ability to resolve projection directionality on its own. By constraining tractography with tracer-informed, blueprint-based direction estimates we obtain a directed scaffold that retains tractography’s subject-specific weighting while adding principled direction information. Furthermore, the method is atlas-agnostic: the blueprint transfer can be applied to alternative parcellations and extended as additional tracer or electrophysiological datasets become available.

We validated the inferred directed scaffold across independent modalities. Edge-level overlap and topology comparisons with cortico-cortical evoked potentials (F-TRACT [35]) reveal substantial convergence between electrophysiological effective connectivity and the DHSC, supporting the biological plausibility of the transferred projections (Fig. 2a,b). At the system level, trophic-level analysis exposes a coherent hierarchical gradient that maps sensory regions to low trophic levels and association and motor regions to higher trophic positions. From a functional perspective (Fig. 4b,c), high-output rather than input nodes demonstrate a distribution that is in alignment with the task-positive region [32] (Fig. 3b). Brain region pairs exhibiting bidirectional structural connectivity demonstrate stronger resting-state functional connectivity in comparison to those exhibiting unidirectional or no connectivity (Fig. 5e), and simulation results that better align with experimental observations during the stimulus-response propagation process (Fig. 7c, Fig. S5). These multimodal validations strengthen confidence of DHSC capturing meaningful aspects of human directed wiring.

### 3.2 Brain network organization based on connectome directionality

Thanks to the incorporation of connection directionality, directed connectomics enables direct inference about the putative flow of information and the large-scale organizational principles of brain networks from static structural architecture. In this study, we leveraged this property by adapting the concept of trophic levels from ecological networks to quantify hierarchical organization in the human brain [27]. This framework provides a complementary and mechanistically grounded perspective on cortical hierarchy that extends prior work based primarily on functional connectivity gradients and microstructural markers.

Over the past two decades, multiple studies using rs-fMRI, intracortical myelin proxies (T1w/T2w), cytoarchitecture, and transcriptomics [52] have converged on a principal macroscale gradient spanning unimodal sensory regions to transmodal association cortex (often termed RSFC gradient-1) [28, 62]. This gradient has been widely interpreted as reflecting a sensory-fugal cortical hierarchy and is frequently used as a proxy for hierarchical organization. However, given the complexity of functional gradients, which emerge from the interplay of multiple factors, the underlying biological significance of these gradients remains unclear. The present DHSC framework enables hierarchical inference directly from directed structural connectivity in the human brain which provides a novel interpretation of functional gradients.

We find that trophic-level–derived hierarchy only weakly correlates with RSFC gradient-1, but shows a substantially stronger association with RSFC gradient-2 (Fig. 4f). RSFC gradient-2 has traditionally been interpreted as reflecting differentiation across sensory modalities. However, emerging work suggests that this gradient may also reflect patterns related to task engagement, dimensionality of representations, and transitions from sensory-driven to integrative and motor-related processing streams [63]. In this context, our findings suggest that RSFC gradient-2 may more directly reflect large-scale feedforward organizational principles, rather than merely modality segregation. From a computational perspective, this interpretation is consistent with architectures of artificial neural networks, in which hierarchical feedforward processing gives rise to layered transformations from input to output representations. The expansion of association cortex in human evolution may therefore be interpreted, in part, as an expansion of intermediate “hidden-layer”–like processing stages, supporting increasingly abstract and integrative computations. Within this framework, trophic levels provide an anatomically grounded metric for such feedforward organization that is not accessible from undirected functional connectivity alone. Despite the network displaying a clear hierarchical structure, the addition of numerous unidirectional edges does not render the network purely feedforward (*F*_0_ = 0.86). Instead, reciprocal and looped motifs dominate the network architecture, consistent with extensive evidence that cortical processing is characterized by dense recurrent and feedback connectivity [20, 64]. This observation underscores that cortical hierarchy is embedded within a strongly recurrent system, in which feedforward structure coexists with pervasive feedback and lateral interactions. This idea also supposed by the results form large-scale simulations, where unimodal signals in the low-trophic level cortex are gradually transformed into multimodal signals in the association cortex which supported by recurrent connections (Fig. 7b, e). Thus, DHSC captures a biologically realistic regime in which hierarchical organization and recurrent dynamics are simultaneously present. In summary, our results demonstrated directed structural connectivity plays a critical role to link classical anatomical hierarchy to functional gradients.

### 3.3 The impact of connectivity directionality on whole-brain dynamics

Finally, we return to the central mechanistic question: how does the introduction of connection directionality, more generally biologically grounded symmetry breaking in large-scale brain networks, reshape whole-brain dynamics, and does this improve the biological realism of physical digital twin models?

Within the model and parameter regime explored here, alterations in network topology, such as the incorporation of long-range connections, primarily enhanced resting-state dynamical fluidity and strengthened global structure–function coupling. This result is consistent with prior large-scale modeling work showing that long-range integration supports metastability and rich spontaneous dynamics. However, since this study focused on the influence of the connectome on whole-brain dynamics (Fig. 5), it examined only the avalanche-critical state in terms of dynamics and did not optimize the functional connectivity, however, the functional connectivity observed following stimulation bore a closer resemblance to the experimental results (Fig. S7). This suggests that ongoing external stimulation, the heterogeneity of brain regions, and E-I balance which are not consider in this study may also be key factors in the brain’s ability to maintain fluidity [65, 66]. Therefore, future research will focus on further optimizing the model, incorporating regional heterogeneity, and exploring the cross-effects of connective directionality and regional dynamics.

However, during stimulus-evoked signal prorogation processing, homogeneous locally dynamic whole-brain model has provided strong evidence that connectome topology and directionality jointly constrain large-scale brain dynamics. This interaction is most evident in the emergence of modular and hierarchical organization in information processing (Fig. 6b). Directionality sharpens functional modularity by segregating rapid, modality-specific responses in primary sensory regions from later, modality-independent activity in higher-order association cortex, consistent with the coexistence of specialized and integrative subsystems [4, 67] (Fig. 6b, 7b). Dynamically,The introduction of directionality provides a structural foundation for transitioning from unimodal rapid responses to modality-independent working memory sustained. The spatial pattern of this sustained activity concentrated in prefrontal cortex and aligns with established observation of distributed working memory [60, 61]. Mechanistically, directionality likely encodes asymmetries in information flow that support efficient feedforward transmission while selectively enabling recurrent processing in higher-hierarchical regions. The findings of Kong et al. also appear to support this point, in that the local dynamical conditions required for resonance differ between the sensory cortex and the association cortex [66]. The sensory cortex displays characteristics that necessitate external electrical input to sustain activity, whereas the association cortex exhibits a stronger capacity for self-activation, at the sametime, these results may also provide a valuable insight into the underlying causes of the current deficits in resting-state functional connectivity. [66]. Together, these results support a hypothesis that directionality governs the transformation and persistence of activity, enabling the coexistence of rapid sensory processing and sustained integrative dynamics in large-scale brain systems. This results also highlight that the connectome featuring directionality and pruning based on biological plausibility may have potential practical value in future computational studies of large-scale whole-brain simulations.

### 3.4 Potential for expansion

However, this framework does not constitute a ground truth human directed connectome. Rather, we provide a principled pathway for inferring directionally informed structural connectivity and an initial, openly accessible prototype of a human directed connectome. It is important to note that this connectome is subject to the following limitations:

First, our method was less accurate in brain regions that expanded greatly during human evolution, particularly within medial temporal and occipital–parietal association cortices (Fig. S2). Because these regions changed so much compared to other species, matching them across species is inherently less reliable. This is a known limitation and an area where future improvements are needed.

Second, threshold selection also remains an open methodological issue. In this study, we focused on a connection density of 20%, selected based on maximal alignment with cortico-cortical evoked potential (CCEP) data and biological plausibility. While this intermediate density achieved an optimal balance between statistical reliability and biological validation in our analyses, prior work has shown that small-world properties and modular organization are preserved across a wide range of network densities in human connectomes. Indeed, we observed robust small-world characteristics across densities from 20% to 60% (Fig. S4), suggesting that many of the qualitative network-level conclusions are not driven by a single arbitrary threshold choice. Nevertheless, systematic optimization of thresholding strategies remains an important topic for future validation.

Finally, the present framework does not yet incorporate the hindbrain including cerebellum, which is increasingly recognized as a critical component of large-scale cortical–subcortical loops and cognitive processing. Extending the directed connectome to include cerebellar circuits, together with targeted experimental validation and cross-species refinement in evolutionarily expanded association cortices, is ongoing essential steps toward improving the completeness and biological fidelity of human directed connectome models.

## 4 Materials and Methods

### 4.1 Inferring directionality from Macaque

#### 4.1.1 Macaque Tracer Data

Modha and Harriger generated the hierarchical directed connectivity matrix for macaques (N = 384 and E = 6,602) was generated by Madha and Harriger in 2010 [18]. In brief, their work integrates over 400 neural tracer papers from the Cocomac database (Modha and Singh, 2010) and organizes the experimentally obtained projections into a binary connectivity matrix, which includes a hierarchical structure. This matrix was then mapped to the D99 brain atlas [31] based on the following rules: (1) If the brain regions share the same name and are described in detail one-to-one, they are directly considered to be the same brain region. (2) If the names of the brain regions are different, then mapping is done based on detailed descriptions or aliases of the brain regions. (3) Integration of multiple brain regions in the higher resolution atlas when the resolution of two more brain regions is inconsistent. The binary connectivity matrix in the D99 atlas (N = 270, cortical region = 158, subcortical regions = 112) was provided in the Supplementary Material.

#### 4.1.2 Tractography data

Tractography data of macaque were taken from open access Brainnetome-8 dataset [12] for building connectome and blueprints. The Brainnetome-8 data set included 2 male and 6 female ex vivo subjects. All subjects consisted of 186 diffusion-weighted images (including 6 non-diffusions for b = 0 *s/mm*^2^, 60 directions for b = 2400 *s/mm*^2^, 120 directions for b = 4800 *s/mm*^2^) with isotropic resolution of 0.3 mm that were acquired using a 9.4 T scanner. For human data, 100 human subjects were randomly selected from the HCP-YA dataset [68].

We utilized the MRtrix3 pipeline [1] to generate a structural connectome-from diffusion MRI (dMRI) data for both human and macaque in this study. For macaque data, Denoising and Gibbs removal were performed. Freesurfers [69] were used to estimate the ranges of white matter (WM), gray matter (GM), and cerebrospinal fluid (CSF) for the estimation of the response function. Based on the segmentation and response function, fiber orientation distributions (FODs) were calculated. Probability tractography was performed using the FODs to generate whole brain streamlines, which are seeded in the white matter mask. For macaque the track length ranges between 2mm to 200mm, each step length is equal to 0.225mm and the maximum curve transfer is 45°, cut off equal to 0.05. Streamline counts and streamline length calculated based on D99 atlas which were registered restarted to structure space using nonlinear registration (SyN) provided by ANTs [70]. For Human subject, all processes for dMRI data follow the HCP pipeline [30]. To generate the connectome, for cortex parcellation follow the HCP-MMP1.0 atlas and for subcortical structure the segmentation was follow the HCPex atlas, which provide 33 subcortical regions per hemisphere. For the cortex part the registration done in the surface, for the subcortical part registration done with in nonlinear registration which is same to macaque.

#### 4.1.3 Cross-species direction mapping

We used the method, Blueprints, to perform cross-species brain region mapping, which is based on protractx2 in FSL [26, 71]. In brief, because macaques and humans share some conservative white matter, we can rely on the relationship between brain regions and white matter tract to evaluate the homology of brain regions between different species.

White matter tracts were generated by Fsl-XTRACT [71]. Each streamline cluster was generated using a set of seed, target, and exclusions mask which provide in standard space (F99 for macaque, MNI152 for human). All processes were performed in the structure space of the subject, and the tractography protocol followed the default setting provided by the XTRACT tool. For cortex structure in both macaques and humans, each vertex of the white-gray matter surface was generated by the freesurfer for probabilistic tractography. The whole brain mask was set as the target. For sub-cortical structures, each voxel in subcortical brain regions were set as seed, whole brain mask set as target. Blueprints were calculated by the Matrix of connections of all voxels in the brain to white matter bundles multiplied by matrix of connections of all vertex/voxels in cortical/subcortical, which characterizes the relationship between brain regions and white matter tracts.

Based on the blueprints the symmetric Kullback-Leibler (KL) divergence was used to measure the similarity of each brain region between macaque and human:

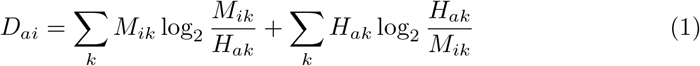

*i* and *a* represent the brain region index in macaque and human. *k* means the white matter tract index.

The distance weighted interpolation is used in mapping the directional connectivity in macaque to humane brain based on the divergence matrix (D). The integration needs performed both in target regions and source regions, thus we infer the projection as follows:

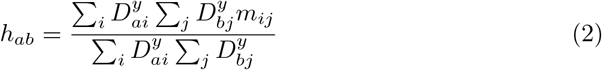

*h*_*ab*_ means the connection between brain regions a, b in human and *m*_*ij*_ means the connection between brain regions i, j in macaque. *y* = -4.

#### 4.1.4 CCEP-based human connectome (F-TRACT, HCP-MMP1.0)

The F-TRACT atlas is based on stereo-EEG (SEEG) electrical stimulation and whole-brain recording in a large cohort of adult patients with pharmaco-resistant focal epilepsy, aggregated across centers (v2307 release; 613 patients, 625 implantations). Connectivity is inferred from statistically significant evoked responses to single-site low-frequency electrical stimulation, providing directed, latency-resolved measures of effective connectivity. Jedynak, M. et al. used the parcellation-based matrices projected onto the MNI-HCP-MMP1.0 atlas (Glasser et al., 2016) provided by F-TRACT. For each ordered pair of parcels (i → j), connectivity is defined as the frequentist probability of observing a significant CCEP response in parcel j following stimulation of parcel i. A response was considered significant when the z-scored averaged evoked signal crossed a threshold of Z = 5 relative to a pre-stimulus baseline. The probability matrix thus represents a directed, weighted connectome reflecting the likelihood of effective signal transmission between parcels. Because F-TRACT data are derived from epilepsy patients and may include indirect (polysynaptic) effects within a 200 ms post-stimulus window, the resulting connectome is interpreted as a probabilistic, effective connectivity map rather than a strictly monosynaptic anatomical connectome.

### 4.2 Graph analysis of directed connectome

#### 4.2.1 Quantification of Small-World Properties Using the Omega (*ω*) Coefficient

The small-world characteristics of the directed network were evaluated using the Omega (*ω*) coefficient [45], a robust metric that simultaneously considers both fundamental properties of small-world networks: high clustering and short path lengths. The calculation procedure is as follows:

First, we computed the actual network’s average clustering coefficient (*C*_*actual*_) and average directed path length (*L*_*actual*_). The clustering coefficient measures the tendency of nodes to form tightly connected groups, while the average path length quantifies the typical separation between any two nodes in the network.

For comparative benchmarking, we generated multiple randomized networks preserving the same number of nodes (N = 426) and identical degree distribution as the original network. From these Erdős-Rényi random networks, we derived the expected average path length (*L*_*random*_). Additionally, we calculated the theoretical clustering coefficient (*C*_*lattice*_) for an equivalent regular lattice network with equivalent connectivity parameters.

The Omega coefficient was then computed using the formula:

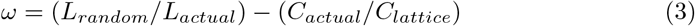

Networks exhibiting small-world properties are characterized by *ω* values approaching zero, indicating they maintain the high clustering of regular lattices (*C*_*actual*_ = *C*_*lattice*_) while achieving the short path lengths of random networks (*L*_*actual*_ = *L*_*random*_). This metric provides a single-valued quantification that effectively discriminates between regular, small-world, and random network topologies.

#### 4.2.2 Trophic level and coherence analysis

To quantify the hierarchical organization and directionality of the human directed connectome, we computed the trophic levels and trophic coherence of the network following the framework proposed by MacKay, Johnson, and Sansom (2020) [27].

Each node *n* in the directed weighted adjacency matrix *W* = [*w*_*mn*_] was assigned: in-weight 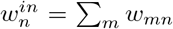 out-weight 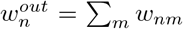 imbalance 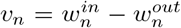 total weight 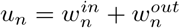.

The (weighted) graph Laplacian operator was defined as:

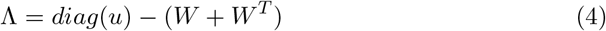

The vector of trophic levels *h* was obtained by solving the linear system:

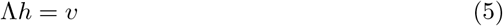

This formulation ensures that networks without basal nodes can still be analyzed.

The degree to which the network edges align along a consistent global direction was quantified by the trophic incoherence:

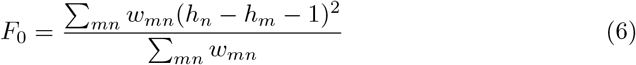

Low *F*_0_ values indicate a strongly hierarchical (coherent) structure, while values approaching 1 correspond to maximally incoherent (balanced or symmetric) networks.

This approach provides a continuous measure of hierarchical flow directionality applicable to any weighted directed network, allowing comparison of structural organization across different brain regions or connectivity models. In this study, we emphasize the network connection pattern, therefore the calculation of trophic level based on the binarized directed connectome. To obtain clear conclusions in subsequent studies, we divided all brain regions into ten groups based on their values of trophic level. Their distribution on the cortex is shown in Figure 2.

#### 4.2.3 Quantification of matrix non-normality

To quantify non-normality of the DHSC, we employed two complementary scalar indices: the Henrici index, which captures global commutator-based non-normality, and a Schur upper-triangular index, which captures modal coupling in the Schur eigenbasis.

Henrici index: The Henrici index was computed from the Frobenius norm of the commutator between *A* and its adjoint:

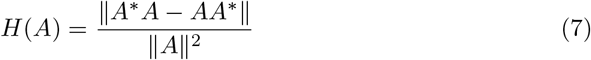

where *A*\* denotes the conjugate transpose of A, and ∥·∥ denotes the Frobenius norm. This normalized index is zero if and only if *A* is normal (*A*\**A* = *AA*\*) and increases monotonically with increasing deviation from normality. The Henrici index thus provides a global, basis-invariant measure of structural non-normality reflecting large-scale asymmetry and directional organization of the network.

Schur upper-triangular index: To quantify modal non-normality, we computed the complex Schur decomposition of *A*:

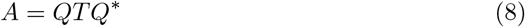

where *Q* is unitary and *T* is upper triangular, with the eigenvalues of *A* on the diagonal of *T* . We extracted the strictly upper-triangular part of *T*,

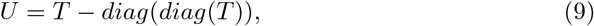

which captures feedforward coupling between Schur modes. The Schur upper-triangular index was then defined as

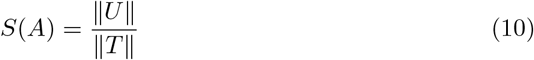

This index is zero for normal matrices, for which *T* is diagonal, and increases with the strength of off-diagonal modal coupling. The Schur index therefore quantifies the degree of eigenmode interaction and provides a dynamical measure of non-normality associated with transient amplification and eigenvector non-orthogonality.

#### 4.2.4 Functional connectivity and its fluidity

For both synthetic and empirical data, we select 9.6-minute time series for calculations.Each regional time series was bandpass-filtered to isolate narrowband fluctuations (0.008–0.08 Hz), since a narrowband signal is required for meaningful phase estimation. The calculation of functional connectivity is based on the Pearson correlation coefficient. The *FC* matrix used for analysis was calculated from the 9.6-minute time series. For functional connectivity fluidity we compute *FC*_*w*_ matrices for each time window *w*, window length is 30s and window step size is 5s. The functional connectivity dynamic *dFC*_*w*_ is calculated as the correlation between different time windows:

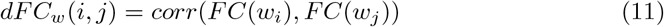

The fluidity of the dynamic functional connectivity *dFC*_*w*_ was defined as the variance of the variance of the values in the upper triangle of the *dFC*_*w*_ with the diagonal offset by the window size.

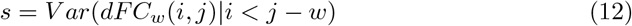

### 4.3 Brain dynamic simulation

#### 4.3.1 Whole brain modeling

We simulate the resting state dynamic in TVB (the virtual brain). Each brain region in the atlas was considered as a node, and its activation was explained by the following function.

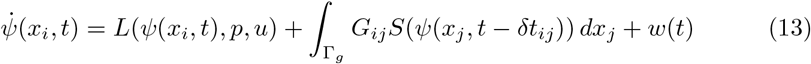

The first term expresses the mean local dynamic of *x*_*i*_ and depends on the local dynamic field *L*, the activation of node *x*_*i*_ at time t *ψ*(*x*_*i*_, *t*), parameters *p* and external input *u*. In this paper, we implement Montbrió model which derived analytically as the limit of infinitely all-to-all coupled QIF (Quadratic Integrate-and-Fire) neurons as the local dynamic for each nodes [56].

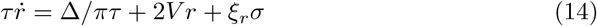

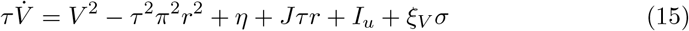

*r*_*i*_ and *V*_*i*_ are the average firing rate and membrane potential of the neural mass, respectively. *η* is the average neuronal excitability; *J* is the synaptic weight inside the neural mass, *δ* is the spread of heterogeneous noise distribution; *τ* is the coefficient used to control the rate of change in a system; *I*_*u*_ is the current input outside the neural mass; *σ* is the noise representing the spontaneous activation of each node, and *ξ*_*r*_ and *ξ*_*V*_ adjust the magnitude of the effect that spontaneous activity has on different variable. To simplify the system and focus on the issues at hand, we configured all nodes with identical parameters to ensure they operate in a bistable state. The settings of the specific parameters are as follows: *τ* = 1.0, *δ* = 0.7, *J* = 14.5, *η* = -4.6, *ξ*_*r*_ = 1.0, *ξ*_*V*_ = 2.0.

The second term expresses the inputs from all connected nodes *x*_*j*_ across the spatial domain Γ_*g*_, which defined by the whole brain network between node *x*_*i*_ and *x*_*j*_, and depends on the connection matrix. The transmission of neural signals from node *x*_*j*_ to node *x*_*i*_ involves temporal delays *G*_*ij*_ and connection strength *δt*_*ij*_ and signal conversion *S* due to synaptic transmission and the propagation of electrical signals along the axon. In this study, *S*(*ψ*) = *G*_*c*_ * *ψ* where *G*_*c*_ is the globular coupling coefficient, *G*_*ij*_ = *log*_10_(*w*_*ij*_ + 1), where *w*_*ij*_ is the streamline counts. *δt*_*ij*_ = *L*_*ij*_*/v, L*_*ij*_ is the streamline length and v = 3.9 m/s.

#### 4.3.2 Resting state simulation

For each synthetic whole-brain time series, we performed a 610-second whole-brain dynamics simulation, the first 10 seconds data were used to complete the transition from the initial state. the hemodynamic response function (HRF) used to compute the BOLD signal, TR = 0.72s.

At the whole-brain macroscale level, we can clearly observe a phase transition from a deactivated (subcritical) to an over-activated state (supercritical), Only at the boundary between these two states (critical boundary) can the connection function, enabling the system to exhibit the fluidity observed in experiments. Hence, we sweep the control parameters *G* and *σ* to find the parameter combination to push the system to the critical boundary. we based on 2 different metric to evaluate

In detail, We first independently sweep 3 different group-averaged connectomes (threshold, directed, directed-symm) in broad range (*G* = [0.01, 0.06], step = 0.0005 ; *σ* = [0.01, 0.06], step = 0.005) to narrow parameter sweep in individual level. individual sweep range were setting as following. threshold: *σ* = 0.05, *G* = [0.030, 0.045], step = 0.00025; Directed: *σ* = 0.045, *G* = [0.040, 0.055], step = 0.00025; Directed-symm: *σ* = 0.05, *G* = [0.030, 0.045], step = 0.00025.

#### 4.3.3 Stimulus perturbations

In this study, stimulation was performed under the resting state. We performed a 80-second whole brain simulation for each stimulation trial, the first 20 seconds were used to complete the transition from the initial state. At 20-seconds, a 2-second square wave was applied to a single brain region to push the brain region into up-state. During the subsequent 60 seconds, we observed the impact of this stimulus on the dynamics of the whole-brain. Since this whole-brain model is noise-driven, its response to stimuli also exhibits randomness. Therefore, we compared responses generated using the same random seed with resting-state activity to obtain its dynamic response. The computational method is as follows:

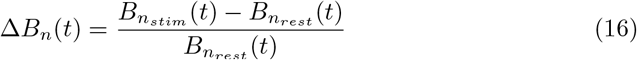

Here Δ*B*_*n*_(*t*) means changes in the BOLD signal at the time point *t*, 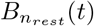 and 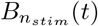 means the observation of BOLD in the resting state and after external perturbations. The simulations of visual and auditory cortex stimulation were repeated 1000 times based on 1000 different random seeds, and the final time series was an average time series aligned based on the stimulation onset times.

### 4.4 Statistics

All neuroimaging data utilized in this study were obtained from publicly available datasets: macaque data from Brainnetome-8 [12] and human data from HCP-YA dataset [29]. A selection of information and key findings will be disseminated on the GitHub platform. The neuroimaging methods, graph theory methods and whole-brain simulation models employed in this study are outlined in the supplementary materials. Statistical analyses were performed in R 4.4.1, and the results are expressed as the mean ± standard error (SEM). All between-group significance tests were conducted using two-tailed unpaired t-tests. Kolmogorov–Smirnov (KS) statistics were used to test for similarity of distributions. Pearson’s correlation coefficient was used for all correlation tests. All statistical results were considered statistically significant at P ¡ 0.05. For correlation tests, a value of ∥ *r*∥ > 0.4 and *P* < 0.05 is considered to indicate a strong correlation.

## Notes

### Competing Interest Statement

The authors have declared no competing interest.

